# On the Contribution of Reproductive and Offspring Investment on Fertility: Human and Animal Societies

**DOI:** 10.1101/2021.05.11.443669

**Authors:** Anuraag Bukkuri

## Abstract

Differences in investment into reproduction or offspring rearing are plentiful throughout the world, from the cells inside our bodies to complex sociological interactions among humans. Such differences can lead to profound impacts on species’ fitness, fertility, and reproductive rates, sometimes in startling ways. In this paper, we create a simple game-theoretical model to *qualitatively* investigate the effects of such differential investment. We focus on fertility in human societies and show that more wealthy individuals produce more offspring within a a mating group. However, when assortative mating mechanisms are introduced, this effectively leads to a speciation event, and a higher reproduction rate for poorer individuals is noticed, capturing what we call the “wealthy-to-poor switch”. We discuss extensions and implications of this work to nupital gifts in ecology and to clonal competition in cancer cell lines under the influence of treatment.

## 1 Introduction

One of the fundamental postulates in sociobiology is that individuals act in ways to maximize their fitness, i.e., to maximize their reproductive potential. Nowhere is this more apparent than in animal ecology, where we see the development of puzzling traits such as the tail of the peacock, the call of a bird, and ritual combat between males for female mating access. All of these traits have withstood the test of time, despite being harmful for the species survival, for the main reason that they aid the organism in gaining more copulations, thereby increasing their fitness. Extreme cases of such phenomena can be seen when one considers altruism and inclusive fitness: worker bees, for example, will forego production of their own offspring to help rear their siblings, with whom they share genes.

It has thus come as a surprise to evolutionary biologists that the demographic transition, a set of stages all human societies progress through as they develop and modernize, seems to have caused a maladaptive outcome for humans [1, 8, 15, 19, 20]. Specifically, it has been noticed that more affluent humans in more developed countries have fewer offspring, on average, than humans in less developed countries. The leading theory in the field is that this phenomena is peculiar to humans and is indeed maladaptive, as those in more developed countries have fewer offspring, thus not acting in a fitness-maximizing manner [1, 6, 10].

However, depending on how we define fitness, perhaps this is not the case. Relative fitness, as contrasted with absolute fitness, is defined as the share an individual has in future gene pools of their *mating group*. Thus, regardless of the size of the mating group, as long as an individual’s genes are represented at a higher than average frequency in the mating group’s gene pool, the individual is reproductively successful [8, 9, 18]. From this, we theorize that individuals in more competitive mating groups must compete with higher quality individuals, thereby changing their reproductive output to align more with that of the group. From a life history theoretic viewpoint, we postulate that individuals in competitive mating groups increasingly shift investment into maintenance from reproduction, leading to a lower absolute fitness but a higher relative fitness. To test these hypotheses, we create a simple game-theoretical model of matings between individuals of different ”classes”. We examine relative reproduction rates, captured by frequencies of phenotypes in the population, of classes of individuals under two conditions: intragroup, random mating and assortative, homophilic mating.

## 2 Model Formulation

To explore these ideas, we first create a simple game theoretical model that captures interactions between poor and wealthy individuals in a population. We shall first focus on basic intra-group dynamics, ignoring the existence of divergent mating groups. Then, we will incorporate a mechanism of assortative mating, and analyze how resulting dynamics differ. The basic pairwise interactions between poor (P) and wealthy (W) individuals are given by the below matrix.

| | $P$ | $W$ |
| --- | --- | --- |
| $P$ | $\frac{M}{2} - c_p$ | $\frac{E(P)}{E(P)+E(W)}M - c_p$ |
| $W$ | $\frac{E(W)}{E(P)+E(W)}M - c_w$ | $\frac{M}{2} - c_w$ |

In the payoff matrix above, *M* represents the payoff from mating. When individuals of the same class interact, they split the payoff in half; when a poor and wealthy individual interact, they split the payoff proportional to the ratio of their wealth, given by *E*(*P*) and *E*(*W*), respectively. In each interaction, the poor and wealthy pay a cost depending on their corresponding investment into their offspring. In the context of human fertility, wealth is captured by the before-tax income of a dual-parent family and the cost is given by the USDA reported average spending per child up until the age of 18.

We can then use the replicator dynamics [2] to probe the dynamics of the frequencies of the poor and wealthy individuals in the mating group. Specifically, if the fitness of the poor and wealthy groups are given as follows:

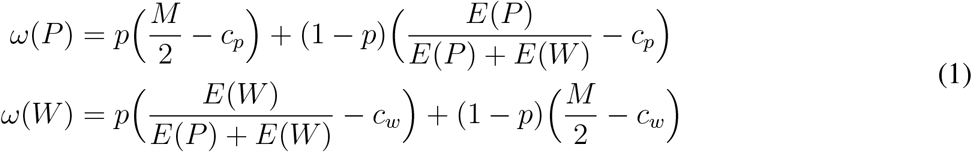

where *p* represents the portion of poor individuals in the population at the given time, the frequency dynamics of the populations can be given as follows:

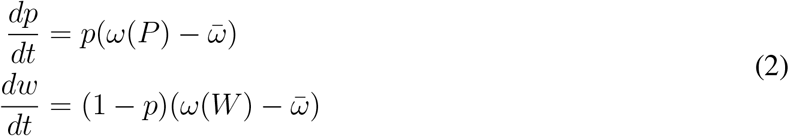

where *w* is the frequency of wealthy individuals and 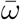, the average fitness of the population, is given by 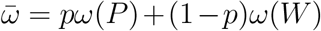. It is worth stating that, though we focus on the case of human fertility here, the model is general enough to apply to any situation in which there is a conflict in a population between investment into reproduction and maintenance, whether by choice or by natural limiting factors such as resource availability. It is important to recognize that the model we present here is only meant to serve as a qualitative model for understanding the underlying dynamics–it is not meant to be used for forecasting population frequencies.

## 3 ESS Analysis

Before we perform some numerical simulations of this model, let’s analyze analyze the evolutionary stable strategies (ESS) of this model. In order for a strategy (in this case, poor or wealthy) to be an ESS, when common in the population, no mutant strategy should be able to invade the population under the influence of natural selection. Stated more formally in our case, this means that for P to be an ESS, one of the following two conditions must hold:

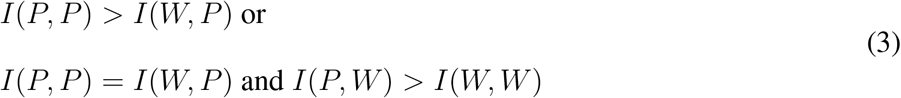

where I(X,Y) denotes the payoff X gets from an interaction with Y. Analogously, in order for the wealthy strategy to be an ESS, we must have:

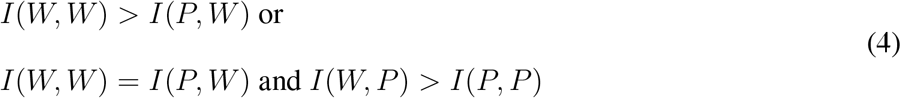

Intuitively, like the well-known Hawk-Dove game, this means that the ESS depends on the balance between the wealth advantage of the wealthy population, and the cost incurred to maintain a wealthy phenotype, *c*_*w*_. If the advantage outweighs the cost, the ESS will one containing all wealthy individuals, but if the cost outweighs the advantage, the ESS will be all poor individuals. In order for a mixed equilibria to exist with the coexistence of both individuals, we require the following two conditions:

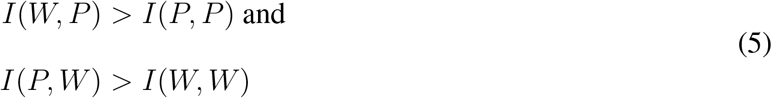

Note that in the case of our model, this is not possible. Thus, we will have either an equilibrium of all wealthy individuals or all poor individuals. The rate at which this occurs provides a marker for the relative reproduction rates of the two phenotypes. For example, if wealthy individuals “invade” the population rapidly, this implies that their reproduction rate is much higher than that of the poor group.

## 4 Random Mating and Intragroup Dynamics

We first consider the case of random mating among members of a mating group. Since we lump all classes of individuals into the same mating group, this means that both ”poor” and ”wealthy” individuals will mate indiscriminately with each other: copulations will only be decided based on frequency of existing populations. Upon first glance, this may not seem like a realistic situation: in human societies, it is quite clear that poor and wealthy individuals do not mate indiscriminately; instead, like mates with like. However, note that here we are interested in *intragroup* mating dynamics, i.e. matings which occur within the *same* mating group. Viewed in this context, the ”poor” and ”wealthy” individuals are most likely part of similar socioeconomically classes. Of course, our assumption of mating groups is a simplification of reality, in which each individual is part of a different mating group and copulations are determined by a myriad of factors apart from socioeconomic background.

To parametrize our model, we use the following data in Table 1, obtained from the USDA annual report: “Expenditures on Children by Families, 2015” [12] and from Statista’s statistics on birth rate by family household income in 2017 [3].

**Table 1:** Values Used to Parameterize our Model

| Characteristic | Lower Class | Middle Class | Upper Class |
| --- | --- | --- | --- |
| Household Income | \$36,300 | \$81,700 | \$185,400 |
| Cost of Child | \$174,690 | \$233,610 | \$372,210 |
| Births/1,000 Women | 57.99 | 51.7 | 45.23 |

The data we use for our model is based on US income, fertility rates, and costs of raising a child solely for data accessibility, conversion simplicity, and to reduce effects of cultural, sociological, or political biases. However, the same ideas, model, and results can be applied to other societies. To parametrize our model, we set E(P) and E(W) to the median household income for the classes under consideration, *M* = 20, and *c*_*p*_ = 3 and multiply this by the ratio in cost of child to obtain *c*_*w*_. Since our model is just meant to qualitatively capture observed trends and not be used as a predictive model, the exact values used for model parameters are not important–the same qualitative results hold for arbitrarily chosen parameter values that preserve class trends in income and birth rate.

We now create three simulations, comparing the upper and lower, upper and middle, and middle and lower class population frequency curves. The results of these simulations can be seen in Figure 1.

**Figure 1:**
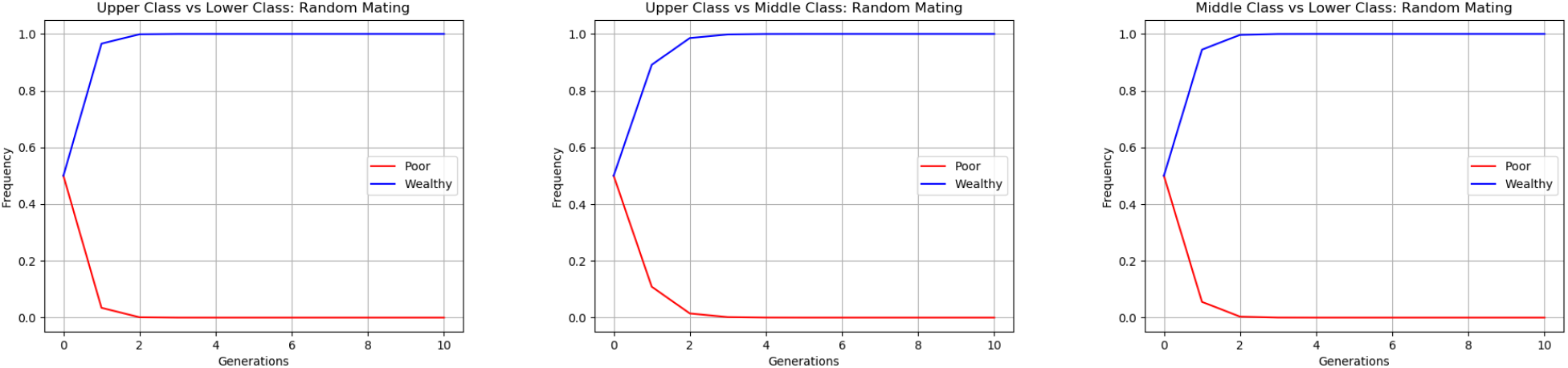
Frequencies of Wealthy and Poor Individuals: Random Mating. Curves were generated by simulating the parametrized replicator equations for 10 time steps (generations). In each case (upper-lower, upper-middle, and middle-lower classes), the wealthy individuals took over the population, pointing to a higher fertility rate for wealthy individuals in intragroup mating systems.

As we can see, in each case, the wealthy population reproduces at a faster rate than the poor population does. This is as expected: in a given mating group, assuming the cost is not exorbitantly high, wealthy individuals will be more attractive mates and will reproduce more often. In fact, this is what we see from the data: the relationship between income and fertility is almost invariably positive within human mating groups [4, 16]. Such studies have also been performed in various primate groups, in which a significant correlation has been noticed between dominance rank and number of mates, measured through displacement and avoidance in mountain gorillas (*Gorilla beringei beringei*) and dyadic pant-grunts in chimpanzees (*Pan troglodytes schweinfurthii*) [21, 22]. But what happens when we introduce mechanisms of assortative mating, thereby creating a divergence in mating groups?

## 5 Assortative Mating and Divergence of Mating Groups

Homophily, the social association of individuals to others similar to them, has been observed extensively through animal and human societies, from social media echo chambers to the formation of dolphin social groups based on foraging techniques and home ranges [13, 14]. Nowhere is this homophily more clear than in mating systems, in which, often, low quality sires and dams mate together and high quality sires and dams mate together. In human marriage systems, the quality of a mate is often determined by socioeconomic standing, captured by before-tax dual household income in our model. This phenomenon has been extensively studied in the sociological and historical literature [5, 11] and is widely recognized as a mechanism for group formation. Note that this assortative mating phenomenon we describe relies on the assumption of self-awareness: that mates can assess their own wealth and the wealth of potential mates. This is fairly common in both animal and human societies, but is an important assumption to keep in mind before extending this model to other contexts. In order to include the effects of assortative mating, we must change the rate at which the poor and wealthy individuals interact among and between each other. To do this, we replace *p* with the term *r* in equation 1 to capture rates of homophily. In this way, we force individuals to interact more frequently with those like them, rather than having their interactions with a given ”class” being simply determined by the proportion of that class in the population. The *r* term can be mathematically defined as follows:

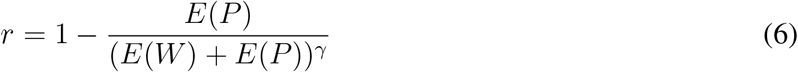

Our formulation assumes that group divergence occurs proportional to the ratios of wealth between the two classes of individuals. Note that when the wealth of the two classes is the same, the groups are entirely overlapping, with *r* = 0.5. As the wealth difference increases, we notice the effects of homophily becoming stronger, with a clearer divergence between groups. The term *γ* is introduced to allow for an additional mechanism for scaling: it represents the rate of inter-group mating that occurs compared to what is expected. Higher values of *γ* correspond to higher levels of segregation between classes, while lower levels of *γ* indicate higher levels of cross-class mating. For the following simulations, we use *γ* = 1.08 to induce greater segregation among classes. With this, we have r = 0.952, 0.912, and 0.904 for the cases of assortative mating between upper and lower, upper and middle, and middle and lower classes. Note that this is in accordance with sociological literature, which indicates that the likelihood of marrying someone of one’s own class varies between classes, with upper classes having a tendency to be more homogamous and the likelihood of homogamy decreasing down the class structure [7, 17].

To compare the prediction from our model to the data, we plot forecasted frequencies of poor and wealthy individuals using the fertility rate data from Table 1. Specifically, letting *f*_*p*_ and *f*_*w*_ represent the fertility rates of the poor and wealthy populations and *p*_*t*_ and *w*_*t*_ represent the frequency of poor and wealthy individuals at generation t, we have:

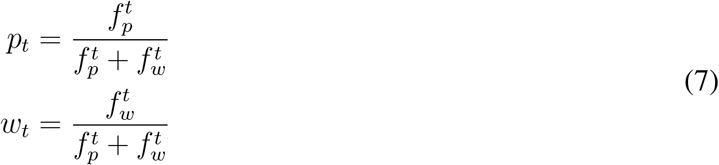

Now, we are ready to perform simulations of the three cases considered before: upper class vs lower class, upper class vs middle class, and middle class vs lower class. The results can be seen in Figure 2

**Figure 2:**
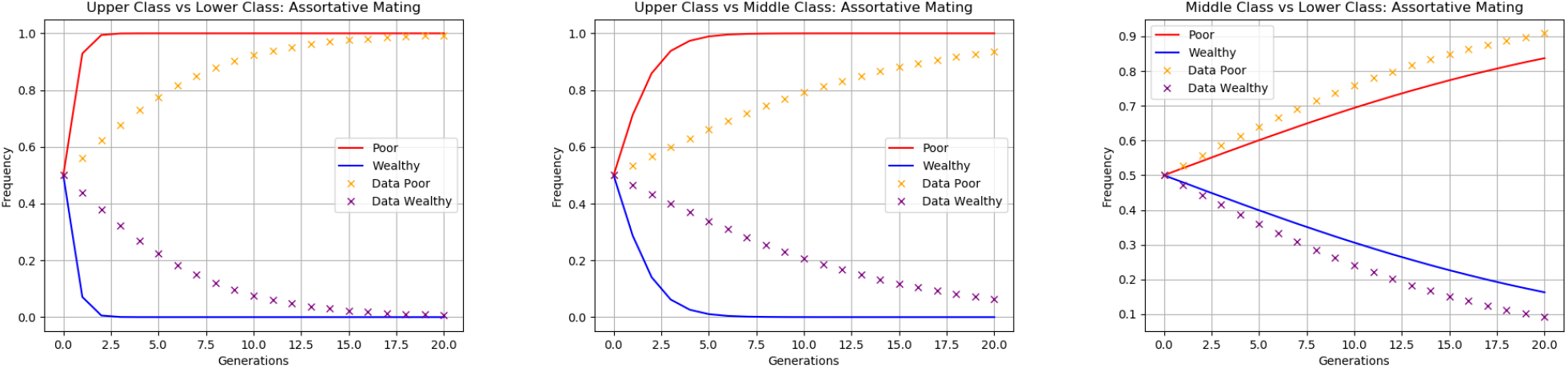
Frequencies of Wealthy and Poor Individuals: Assortative Mating. Curves were produced by simulating the parameterized replicator equations, with the assortative mating term, for 20 time steps (generations). Forecasted data (denoted by x) was plotted by using equation 7 with the relevant fertility data from Table 1. By incorporating the assortative mating mechanism, in all three cases, we capture the “wealthy-to-poor shift” in which poor individuals have higher fertility rates.

As we can see, in all cases, the model captures the “wealthy-to-poor switch”: it predicts a higher frequency (and thus reproduction rate) of poor individuals than wealthy individuals, in accordance with the data. Though our model does not fit the data particularly well, it does qualitatively reproduce the observed trends in fertility. When assortative mating was introduced into the system, effectively creating a quasi-speciation event, we notice the emergence of r- and K-like phenotypes. The poorer individuals act like r-selected species: they are of low quality but reproduce frequently, while the wealthy individuals are K-selected, producing few, high-quality offspring. Interpreting this in a life history-theoretic point of view, the poor individuals tend to invest more in reproduction and less in maintenance than their wealthier counterparts. Thus, our *qualitative* model, in accordance with the data, proposes that, within mating groups, more wealthy individuals have higher rates of reproduction. However, a tragedy of the commons phenomenon arises when assortative mating is introduced. In more competitive mating groups, in order to maintain a high *relative* fitness, individuals sacrifice *absolute* fitness. Thus, we notice that individuals in less competitive mating groups have higher absolute reproductive rates than those in more competitive groups.

## 6 Conclusion

In this paper, we have used a simple game-theoretical model in an attempt to reconcile the fertility paradox in sociobiology: why more affluent individuals tend to have fewer children than less affluent individuals. Though our model solely replicates qualitative trends in fertility rates and is simplistic in that it ignores cultural factors, it is able to reproduce the ”wealthy-to-poor” switch through the introduction of an assortative mating mechanism. Namely, we have shown that, within mating groups, wealthy individuals have a higher fertility rate. However, when mating group divergence is permitted, wealthy individuals are placed into more competitive mating groups and their fertility rates are markedly lower than those of poor individuals.

Though we focus on the case of human fertility here, this work has much broader implications, applying to any situations where there is conflict in a population between investment into reproduction and maintenance. For example, the same model can be used to investigate nuptial gift presentation in various species, from great grey shrikes to arachnids. Namely, it can help us understand how the quality of competing males and the target female is related to the quality of gift that is presented. As another example, the model can be extended to investigate clonal competition among sensitive and resistant cells under treatment, in which quality is encoded as treatment resistance and cost represents the cost of resistance.

